# ModBERT: a DNA modification and genetic variant-aware DNA large language model

**DOI:** 10.64898/2026.11.12.766258

**Authors:** Dante Bolzan, Ramana V. Davuluri, Shaun Mahony, Ferhat Ay

**Affiliations:** La Jolla Institute for Immunology, La Jolla, CA, 92037, USA; Bioinformatics and Systems Biology PhD Program, University of California, San Diego, CA, USA; Department of Biomedical Informatics, Stony Brook University, NY 11794, USA; Center for Eukaryotic Gene Regulation, Department of Biochemistry and Molecular Biology, The Pennsylvania State University, University Park, PA 16802, USA; Department of Pediatrics, University of California San Diego, La Jolla, CA, 92093, USA

## Abstract

The development of foundation models that learn a general grammar of DNA (DNA language models) has accelerated in recent years. The vast majority of DNA large language models (LLMs), such as DNABERT-2 and Nucleotide Transformer, are pretrained almost exclusively on reference genomes. They therefore cannot represent the two sources of variation most directly tied to regulatory differences between people: genetic variants and DNA methylation. Long-read sequencing (Nanopore and PacBio) now measures both from the same native DNA molecules, but no model is designed to use this joint readout. With this in mind, we developed ModBERT, a DNA language model built on a five-letter probabilistic genome representation in which each position is a distribution over A, C, G, T, 5mC (represented with M). In pretraining, these distributions encode population allele frequencies and CpG methylation levels from 242 individuals in the 1000 Genomes Long-Read Sequencing Consortium, learned with a soft-target masked language modeling objective. We further propose probabilistic fine-tuning, in which the same representation encodes one individual’s genotype and methylation. A fine-tuned model can then make person-specific predictions from a single assay. Under this novel fine-tuning framework, we show that ModBERT effectively utilizes methylation information to classify peaks from functional genomic assays, including histone mark (n=11 different marks) and transcription factor binding (n=69 TFs) ChIP-seq data, outperforming ab initio trained models and ModBERT pretrained on sequences without methylation. In silico methylation perturbations yield TF sensitivities that correlate with independent methylation-aware motif models. Using an evaluation design that separates generalization to new loci from generalization to new individuals, ModBERT predicts chromatin accessibility at held-out loci in held-out individuals (AUROC 0.88). Its predicted variant effects agree with likely causal caQTLs (Spearman ρ = 0.52) but not with variants associated only through linkage, a traditionally challenging task. Overall, ModBERT establishes a framework for personal-genome language modeling, in which a model reads each individual’s genetic and epigenetic sequence rather than a shared reference. This enables *in silico* dissection of variant and methylation effects on gene regulation and principled prioritization of candidates for experimental validation as population-scale long-read cohorts (e.g., 1KGP and Human Pangenome Reference Consortium) grow.

## INTRODUCTION

The development of DNA Large Language Models (LLMs) that learn a general grammar of DNA has accelerated in recent years. These models are pretrained on reference genomes of one or more species. The pre-training phase may employ either masked language modeling (MLM) or causal language modeling (CLM) with the type of tokenization being either single DNA bases (A, C, G, T), k-mer (DNA sequence of length k), or BPE. The fine-tuning stage consists of sequence or token-level tasks such as predicting whether a sequence contains a regulatory element or variant effect prediction. The vast majority of existing DNA LLMs such as DNABERT^1^, DNABERT-2^2^, Nucleotide Transformer^3^, and HyenaDNA^4^ are pre-trained on reference genomes, ignoring the critical regulatory information conveyed by DNA modifications and genetic variation.

Single Nucleotide Polymorphisms (SNPs) are widely-studied genetic variants associated with disease risk, complex traits, and drug response. 5-methylcytosine (5mC), known as a “fifth base”, is a primary epigenetic mark typically associated with transcriptional repression. DNA methylation is implicated in cellular differentiation, gene regulation, and genomic stability^5^. Aberrant DNA methylation patterns have been widely implicated in aging and in numerous complex diseases. The mechanisms that govern site-specific methylation remain incompletely understood. Evidence suggests that flanking nucleotide sequences and CpG sites contribute to the likelihood of CpG methylation^6^. MethylQUEEN^7^ utilizes DNA methylation information during pretraining by representing 5mC as an “M” whereas MutBERT^8^ incorporates SNP information by representing each SNP as a probability distribution over A, T, C, G bases based on their allele frequencies in the human population.

Here, we developed ModBERT, a DNA LLM for the human genome that incorporates both SNPs and DNA methylation (5mC) during pretraining with the goal of learning representations that better capture functional genomic contexts and human population variance.

## RESULTS

### ModBERT pretraining and fine-tuning frameworks

To first learn representations that capture general genomic and epigenomic grammar, ModBERT was pretrained on 512-bp length sequences derived from the human reference genome overlaid with population-level SNP allele frequencies and CpG methylation percentages derived from lymphoblastoid cell lines (LCLs) of a diverse cohort of 242 individuals sequenced using Oxford Nanopore technology by the 1000 Genomes Long Read Sequencing Consortium (1KGP-LRSC)^9,10^. These population-level frequencies were encoded by representing input sequences as probabilistic matrices with the model trained to reconstruct probabilistic vectors over the four canonical bases (A,C,G,T) and 5mC (M) of masked sites in the sequence **(Figure 1a**). The probabilistic representation of sequence and ModBERT’s architecture allow the model to learn nucleotide-level embeddings that capture both genetic variation and methylation frequencies genome-wide (**Methods**).

**Figure 1:**
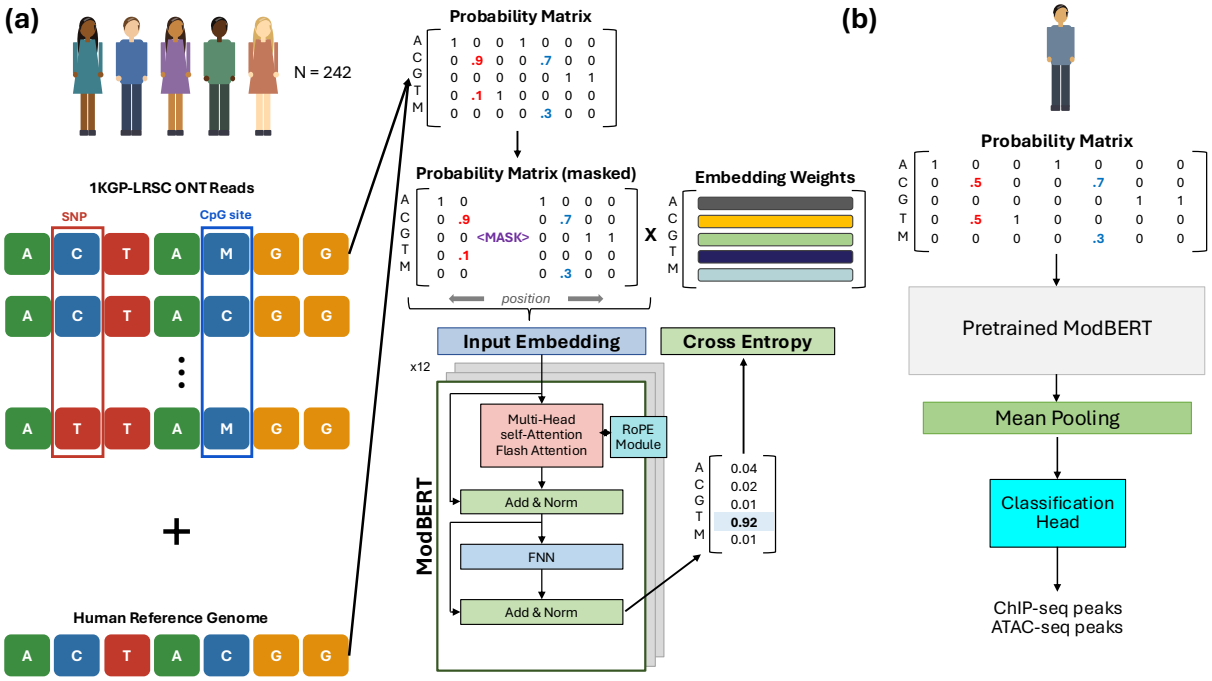
Overview of pretraining and fine-tuning. **(a)** CpG site and SNP identification with accompanying methylation fraction and allele-frequencies respectively derived from Oxford Nanopore DNA sequencing of 242 individuals in the 1000 Genomes Long Read Sequencing Consortium (1KGP-LRSC). ModBERT performs masked language modeling at single-nucleotide resolution. **(b)** Probabilistic fine-tuning on individual samples. Pre-trained model’s weights are updated along with classification head.

Building on these pretrained representations, we fine-tuned ModBERT on individual-level labeled tasks. In lieu of standard fine-tuning in the DNA language modeling field on reference genome sequences, we developed a novel probabilistic fine-tuning approach on data from specific individuals (**Figure 1b**). In this formulation probabilistic vectors representing single nucleotide variants directly encode an individual’s zygosity while CpG site vectors encode an individual’s fraction of reads calling it as methylated. The probabilistic matrix is then fed into pretrained ModBERT to obtain nucleotide-level embeddings. These embeddings are then aggregated via mean pooling and fed into a linear classification head for binary ChIP-seq and ATAC-seq peak identification. During fine-tuning ModBERT weights are updated jointly with the task-specific head, allowing the pretrained representations to adapt fully to each downstream task. Probabilistic fine-tuning presents the model with an individual’s variants and methylation levels, all of which are linked to transcription factor binding, chromatin accessibility, and histone modifications. This allows ModBERT to make individual-specific predictions and to learn how genetic and epigenetic variation along with reference sequences jointly contribute to regulatory outcomes. **ModBERT’s pretrained sequence-level embeddings capture genomic elements and methylation sensitivity**

Before evaluating ModBERT on fine-tuning tasks, we conducted zero-shot tasks to assess whether ModBERT’s pretrained sequence embeddings already distinguish different genomic elements. We sampled 5,000 512-bp length sequences from each of five genomic elements genome-wide. The sequences, represented probabilistically in the same fashion as the sequences used for pretraining ModBERT, were then fed into 0the pretrained ModBERT model to obtain sequence-level embeddings from each of the model’s 12 layers. UMAP projections of these embeddings reveal that ModBERT can already distinguish genomic elements, with separation of 5’ UTR sequences beginning in layer 2 and marked separation of coding sequences and a subset of intergenic sequences beginning in layer 7 (**Figure 2a**). The lack of separation for 3’ UTR sequences corroborates zero-shot evaluations of Nucleotide Transformer^3^, suggesting that the incorporation of genetic and epigenetic variation during pretraining does not aid in the distinction of this element type. We also pretrained ModBERT on sequences without 5mC (setting M = 0 for all CpG sites) to assess the impact of methylation on pretrained representations. After feeding this model the same set of sequences overlaid with SNP allele frequencies but lacking 5mC (since the model did not learn an “M” embedding during pretraining), we found that the 5’ UTR sequences do not separate as well as the full model embeddings (**Figure 2b**). We also observed slightly less separation of coding sequences compared to the original ModBERT embeddings for layers 7 and 12. This is consistent with methylation levels being higher in exons compared to introns, independent of GC-content differences^11^.

**Figure 2:**
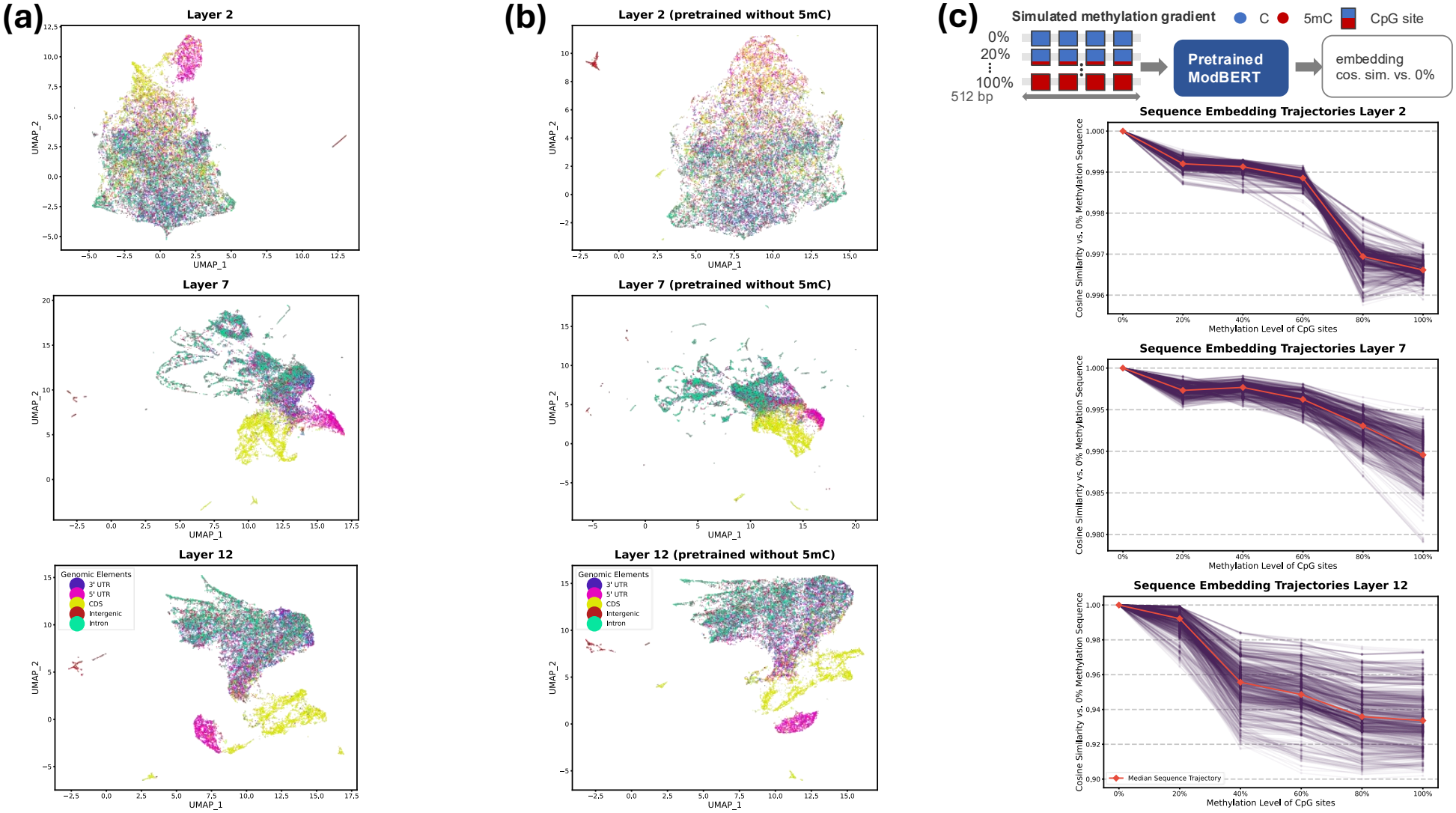
ModBERT sequence embeddings capture genomic elements and exhibit methylation sensitivity. **(a)** UMAP projections of ModBERT sequence embeddings extracted from specific layers. CDS = coding sequence. **(b)** same as (a) but here ModBERT was pretrained on sequences without 5mC and fed the same sequences without 5mC. **(c)** Simulated methylation gradient where each sequence’s CpG sites are uniformly set to varying methylation percentages. Spaghetti plots of cosine similarity of each methylated sequence to its 0% methylated baseline. Each line corresponds to a unique sequence. Each sequence contains the same number of CpG sites.

To assess more directly how ModBERT incorporates methylation into its pretrained representations, we simulated a methylation gradient across 1,000 512-bp sequences randomly sampled from autosomal chromosomes (**Figure 2c**). All sequences contain exactly 30 CpG sites. For each sequence, we generated six versions in which the methylation level of every CpG site was set uniformly to 0%, 20%, 40%, 60%, 80%, or 100%, and computed the cosine similarity between the sequence embedding of each version and that of its unmethylated (0%) counterpart at each layer of the model. Across all layers the median cosine similarity across sequences tends to drop as methylation increases with greater drops as embeddings are extracted deeper in the model (median cosine similarity of ∼0.997 for full methylation at layer 2 while dropping to ∼0.99 and ∼0.93 for layers 7 and 12, respectively). While this monotonic decrease suggests that ModBERT encodes methylation level as a graded rather than binary signal, different layers exhibit different median trajectory shapes, with some degree of a threshold-like pattern at different methylation levels. The steepest decline in median cosine similarity occurs between 60% and 80% in layer 2 and between 20% and 40% in layer 12, while layer 7 exhibits a more gradual decline over all methylation levels. Heterogeneity in the sequence trajectories increases in deeper layers, suggesting that the effect of methylation on ModBERT’s representations depends on sequence context. All in all, these results indicate that ModBERT progressively integrates methylation with sequence information across its layers, consistent with the improved separation of genomic elements observed in intermediate and late layers.

### ModBERT accurately predicts histone ChIP-seq peaks in GM12878

Given that ModBERT’s pretrained embeddings already distinguish genomic elements and respond to methylation in a context-dependent manner, we next performed probabilistic fine-tuning on ModBERT for sequence classification tasks using ENCODE^12^ ChIP-seq data from GM12878, a cell line not present in the pretraining sample set, but for which we obtained ONT data (**Methods**). We first assessed ModBERT’s ability to classify ChIP-seq peaks in eleven different histone marks. Positive sequences were constructed by extracting 1,001-bp length sequences from the center of ChIP-seq peaks, while GC-content matched negative sequences of equal length were extracted from regions not overlapping any peaks (**Figure 3a**). Training and testing were done independently for each histone mark. ModBERT achieved a mean accuracy of 86% across the histone marks, outperforming non-pretrained (ab initio) baseline models as well as the ModBERT model pretrained on sequences lacking methylation information (ModBERT_no_5mC) (**Figure 3b**). Notably, ModBERT achieved a higher accuracy than ModBERT_no_5mC for each histone mark, demonstrating that incorporating 5mC during pretraining helps with general histone mark peak identification.

**Figure 3:**
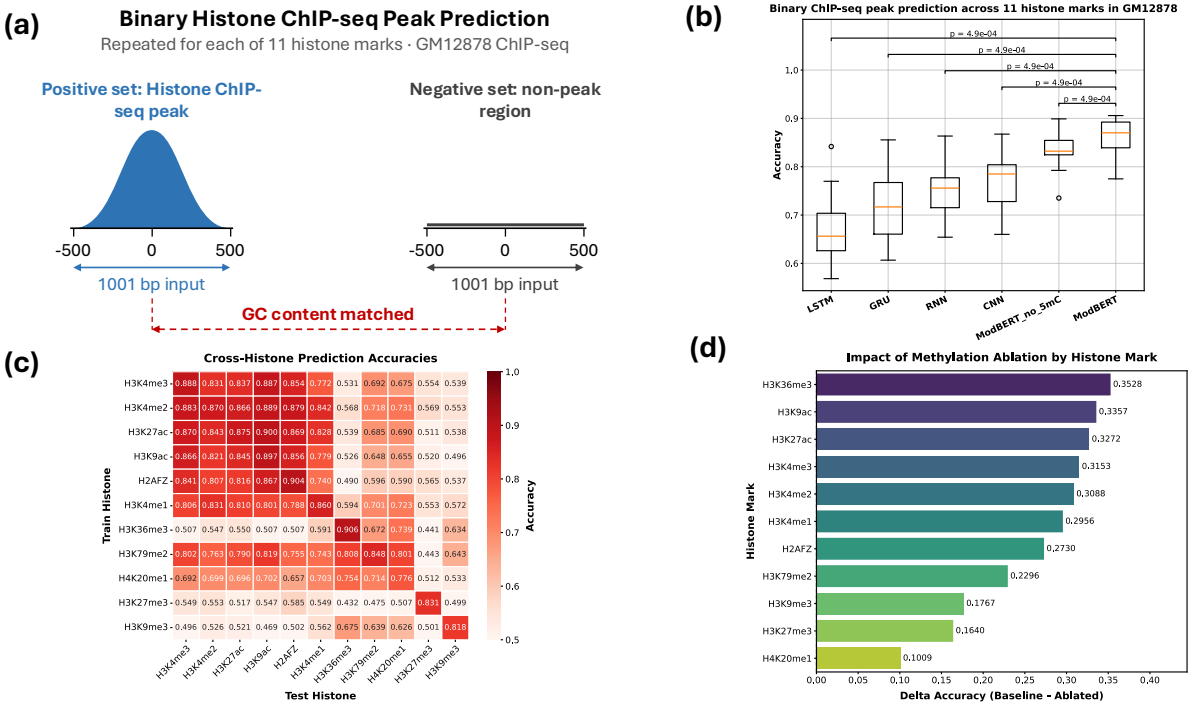
Binary histone ChIP-seq peak prediction across 11 histone marks in GM12878. **(a)** Positive and negative set construction. **(b)** Accuracy in left out chromosomes 5, 10, 14, 18, 20, 22. Each data point corresponds to a separate histone mark. The Long Short Term Memory (LSTM), Gated Recurrent Unit (GRU), Recurrent Neural Network (RNN), Convolutional Neural Network (CNN) models were trained ab initio. ModBERT_no_5mC is pretrained on sequences without 5mC. P-values are from one-sided Wilcoxon signed-rank tests for ModBERT with respect to other models. **(c)** ModBERT cross-histone results from taking the trained fine-tuned model of each histone mark and testing it on all other histone marks. **(d)** ModBERT delta accuracies from removing methylation in test positive and negative sequences. Ablation accuracies are computed over the same positive and negative sequences as the baseline accuracies but with 5mC probabilities set to 0 for all CpG sites.

In addition, cross-histone mark testing of ModBERT’s fine-tuned models captured known chromatin state relationships (**Figure 3c**). Models trained on marks associated with active promoters and enhancers (H3K4me3, H3K4me2, H3K27ac, H3K9ac, H2AFZ, and H3K4me1) generalized well to one another, whereas models trained on marks of transcribed gene bodies (H3K36me3, H3K79me2, H4K20me1) transferred primarily within that group, with H3K79me2, which peaks just downstream of the transcription start site (TSS) and is found in some enhancers, bridging both groups^13,14^. Models trained on repressive marks (H3K27me3 and H3K9me3)^15^ were largely specific to their own mark, and the H3K27me3 model performed at or slightly below chance on transcribed gene body marks, most strongly for H3K36me3 (0.432), consistent with these two marks being largely mutually exclusive^16,17^.

We next performed a methylation ablation experiment to assess the fine-tuned ModBERT models’ sensitivity to 5mC when distinguishing between positive and negative sequences. We set the methylation probability to zero at all CpG sites in both positive and negative sequences in the test set and fed these 5mC-less sequences into the trained fine-tuned models. This ablation led to a drop in accuracy in all histone marks, with the magnitude of reduction closely following the chromatin-state groupings captured in the cross-histone analysis (**Figure 3d**). The largest decreases occurred for active promoter and enhancer marks (H3K9ac, H3K27ac, H3K4me3, H3K4me2, and H3K4me1; decreases of 0.30–0.35), reducing these models to near-chance accuracy. Intermediate decreases were observed for H2AFZ (0.27) and H3K79me2 (0.23), whereas the repressive marks H3K9me3 (0.18) and H3K27me3 (0.16) showed smaller decreases, and H4K20me1 was least affected (0.10). This ordering is consistent with the distinctive methylation landscape of active regulatory elements, which are characteristically hypomethylated relative to the surrounding genome, making methylation a highly discriminative feature for these marks. H3K36me3, which marks transcribed gene bodies, also showed a large decrease upon methylation ablation (**Figure 3d**). Although gene bodies are generally methylated, this broad methylation pattern may itself serve as a discriminative feature distinguishing H3K36me3 peaks from GC-content matched background. In contrast, H4K20me1 was least affected, suggesting that for this mark the model relies primarily on sequence features rather than methylation. These results also mirror the pattern of improvement of ModBERT over ModBERT_no_5mC, where the largest gains occurred for active marks.

Given that GC content does not specifically control for CpG frequency, which tends to be higher in active regions, it is conceivable that a model could learn to separate positives from negatives from differential CpG frequencies instead of learning to utilize 5mC levels. To test this, we repeated all experiments using dinucleotide-matched, real genomic sequences found outside peaks as the negative set. ModBERT still achieved a significantly higher accuracy than the other models, suggesting that at least some of its competitive edge is due to learned 5mC grammar during pretraining rather than CpG density patterns (**Supplementary Figure 1b**). Together, these results show that pretraining with 5mC yields representations that not only improve histone mark peak classification but also capture biologically meaningful relationships between chromatin states.

### ModBERT accurately predicts transcription factor ChIP-seq peaks in GM12878

Having established that ModBERT captures chromatin-state relationships across histone marks, we next asked whether it could resolve the finer-scale, sequence-specific binding of transcription factors (TFs), using ChIP-seq data for 69 TFs in GM12878. Here, we extracted a 101-bp sequence centered in peaks for the positive set with GC-content matched negatives obtained from non-peak background as in the histone prediction task (**Figure 4a**). ModBERT again achieved a higher mean accuracy (86%) than the ab initio models (70%-84%) (**Figure 4b**). ModBERT shows a slight, statistically significant improvement over ModBERT_no_5mC with a higher accuracy in 50 of the 69 TFs, suggesting 5mC variation during pretraining helps with discriminating TF bound sites, consistent with the known methylation sensitivity of some TFs^18^. To more systematically probe the methylation sensitivity of the fine-tuned TF ModBERT models, we fully methylated and demethylated (artificially set methylation to 100% and 0% respectively at all CpG sites) the positive sequences in the test set for each TF. We then fed these perturbed sequences into each fine-tuned model to compute a mean delta predicted binding probability (full methylation - full demethylation) across all positive sequences. We found that ModBERT deemed all 69 TFs to preferentially bind to unmethylated sequence (delta < 0), with delta values ranging from -0.92 to -0.02. To validate these findings, we computed methylation sensitivity scores derived from the methylation-aware motif models from Grau et al^19^. These models are GM12878-specific and were available for 59 of the 69 TFs we investigated. To more confidently assess methylation sensitivity, we excluded 15 TFs that lacked prominent CpGs in their predicted binding sites (**Methods**). Negative sensitivity scores indicate preferential binding to unmethylated DNA. The Grau et al. models agreed with ModBERT in predicting binding antagonism to methylation for all TFs with a modest correlation between ModBERT’s delta binding probabilities and the sensitivity scores (**Figure 4c**). ModBERT largely captures TF-specific differences in methylation sensitivity without being given motif annotations.

**Figure 4:**
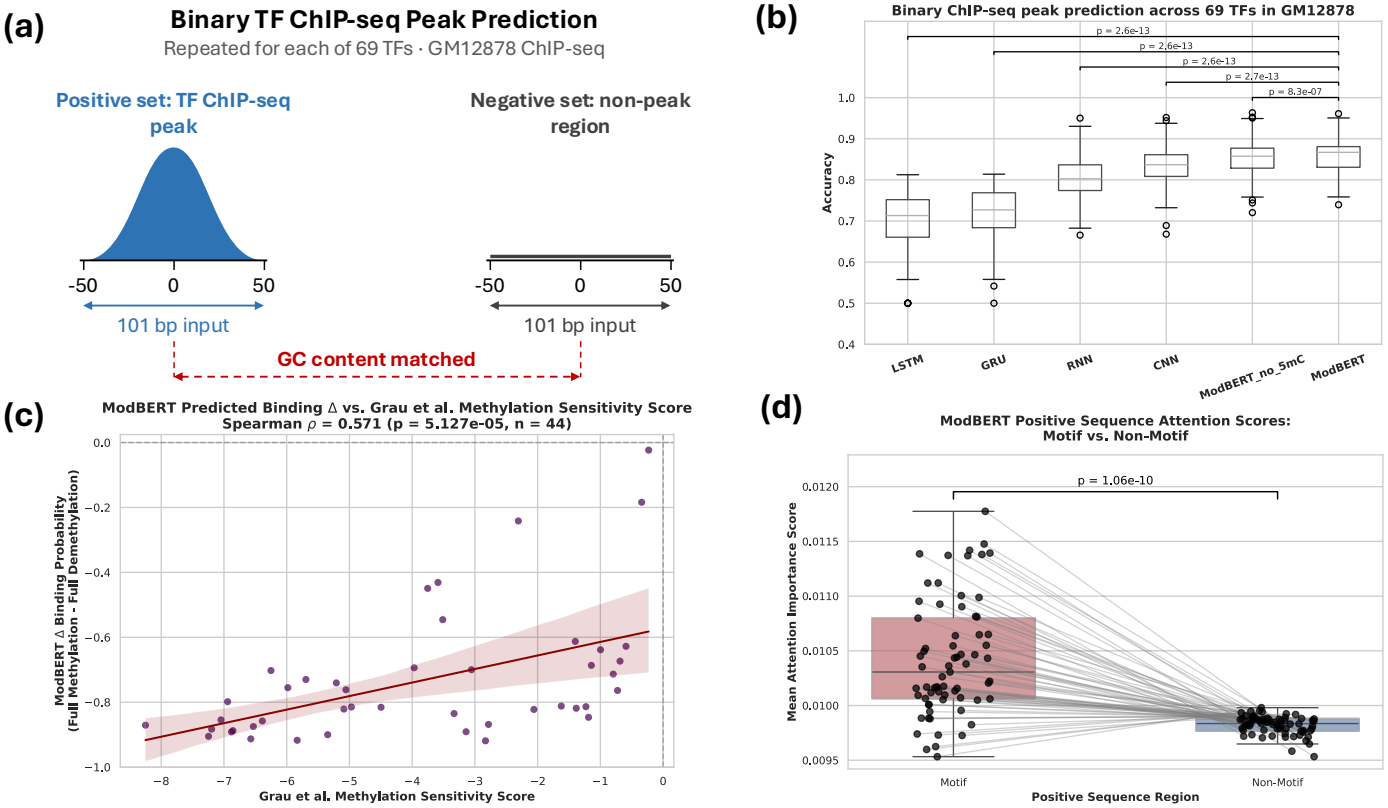
Binary TF ChIP-seq peak prediction across 69 TFs. **(a)** Positive and negative set construction. **(b)** Accuracy in left out chromosomes 5, 10, 14, 18, 20, Each data point corresponds to a separate TF. P-values are from one-sided Wilcoxon signed-rank tests for ModBERT with respect to other models. **(c)** Scatterplot of ModBERT predicted delta binding probability and Grau et al. methylation sensitivity score for 44 out of the 69 TFs. Each data point corresponds to a TF. Note that the predicted delta is a mean across all positive sequences. **(d)** Mean attention score between motif and non-motif regions in positive sequences. Each dot corresponds to a TF with lines connecting motif and non-motif mean attention scores across positive sequences of the same TF. P-value is from a one-sided Wilcoxon signed-rank test.

Lastly, to examine whether ModBERT attends to TF binding motifs, we compared mean attention scores at motif and non-motif positions within positive sequences (**Figure 4d**). Motif instances were identified using FIMO^20^ with HOCOMOCO v14^21^ motifs. For most TFs, motif positions received higher mean attention than flanking positions, indicating that the fine-tuned models focus on the sequence features known to mediate TF binding although the differences were small. These results show that ModBERT accurately predicts TF binding, attends preferentially to the motifs that mediate it, and captures TF-specific methylation sensitivity consistent with independent motif-based estimates.

### ModBERT accurately predicts ATAC-seq peaks within and across individuals

Having shown that ModBERT can predict regulatory features within a single cell line, we next evaluated its capacity to predict chromatin accessibility across individuals (different donor samples), by fine-tuning it on ATAC-seq data from lymphoblastoid cell lines from 27 individuals profiled in Kumasaka et al.^22^ and which were also in the 1KGP-LRSC 242-sample cohort, with each individual’s sequences represented using their own genotypes and methylation levels. We formulate ATAC-seq peak prediction as a binary classification problem using 201-bp windows centered on peaks, following the same positive and negative set construction framework as in the GM12878 tasks. In addition to the unseen chromosomes (peaks) task that was employed with GM12878 ChIP-seq peak predictions, here we also carried out an unseen samples and an unseen samples and peaks task (**Figure 5a**). In the unseen samples task, models were trained on a subset of individuals and tested on the remaining five individuals across all chromosomes. In the unseen samples and peaks task, the most stringent task, models were tested on held-out chromosomes in held-out individuals, so that neither the test peaks nor the test individuals were seen during training.

**Figure 5:**
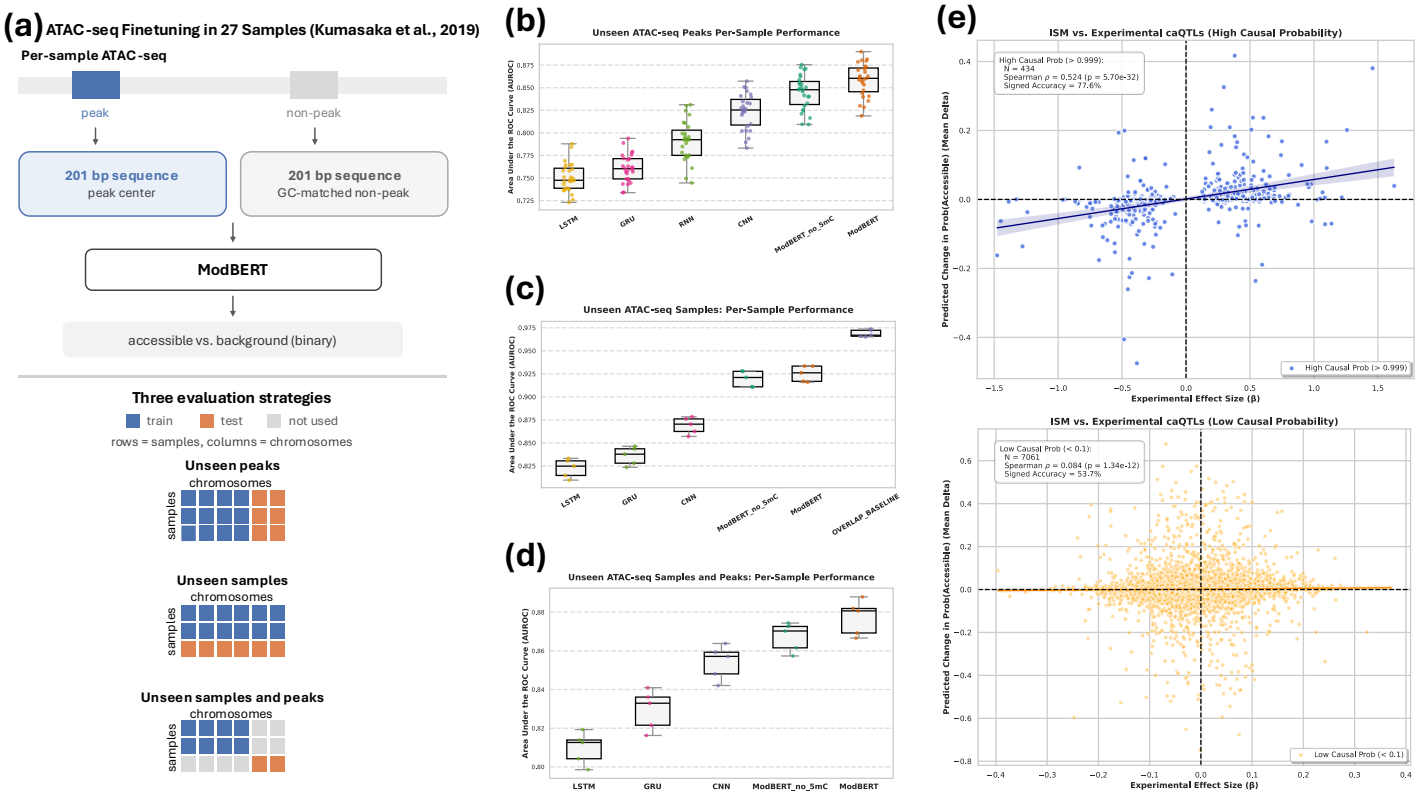
ATAC-seq fine-tuning in 27 samples (individuals). **(a)** Training and testing schematic. Chromosomes 5, 10, 14, 18, 20, 22 were left out whenever peaks were unseen. The same five samples were left out in both the unseen samples and unseen samples and peaks task. **(b)-(d)** Per-Sample AUROC for each of the three evaluation strategies. Note that the RNN model is not included in (c) and (d) due to divergence during training. **(e)** In Silico Mutagenesis (ISM) by one-hot encoding the reference (REF) and alternate (ALT) allele of caQTLs found in unseen peaks in unseen samples and computing a delta = Prob(ALT) - Prob(REF) per sample using the unseen samples and peaks model. The scatter plots plot the mean delta across samples as a function of the experimental effect size (β) (each dot is a caQTL-peak pair). The two subplots are stratified by high and low causal probability, a metric utilized by Kumasaka et al. to determine the probability of a variant being the causal caQTL of a peak.

ModBERT achieved the highest per-individual performance among the deep learning models in all three settings. For unseen peaks, ModBERT reached a mean AUROC of 0.86 across samples, compared with 0.85 for ModBERT_no_5mC and 0.82 for the best ab initio model (CNN) (**Figure 5b**). For unseen samples, ModBERT achieved a mean AUROC of 0.93, compared with 0.92 for ModBERT_no_5mC and 0.87 for the CNN (**Figure 5c**). In this task, however, a simple overlap baseline that predicts the accessibility of a sequence in a held-out sample based on the number of training samples that contain a peak that overlaps with it achieved a higher AUROC (0.97), indicating that most accessible regions are shared across individuals and that a model can perform well here largely by recognizing/memorizing regions accessible in other samples. The unseen samples and peaks setting removes this advantage, since the test regions were never observed during training. Here, ModBERT again performed best, with a mean AUROC of 0.88, compared with 0.87 for ModBERT_no_5mC and 0.85 for the CNN (**Figure 5d**), showing that its performance on new individuals does not depend on having seen the same regions during training. Across all tasks, 5mC during pretraining only slightly improved performance in the fine-tuning.

Because accessibility is largely shared across individuals, aggregate classification metrics provide limited insight into whether ModBERT captures differences between individuals. To test this directly, we compared ModBERT’s predicted variant effects with lead chromatin accessibility quantitative trait loci (caQTLs) identified by Kumasaka et al. We focused on lead caQTLs found within the 201-bp length positive sequences in unseen peaks in unseen samples. For each caQTL and sample, we performed an in silico mutagenesis (ISM) by substituting the variant with a one-hot encoding of the reference allele and the alternate allele and computing a mean delta in predicted accessibility across unseen samples using the unseen samples and peaks model. Among variants with high probability of being causal (as determined by Kumasaka et al.’s pairwise hierarchical model), predicted effects correlated with experimental effect sizes (Spearman ρ = 0.524, p = 5.7 × 10^−32^), and ModBERT predicted the correct direction of effect for 77.6% of variants (**Figure 5e**). In contrast, for variants with low causal probability, which are likely associated with accessibility only through linkage disequilibrium with causal variants, the correlation was weak (Spearman ρ = 0.084) and directional accuracy was near random (53.7%). ModBERT’s signed accuracy also increases as a function of the causal probability threshold (**Supplementary Figure 3**). These results indicate that ModBERT’s predicted effects reflect the regulatory consequences of the variants themselves rather than their statistical association with accessibility through linkage. Interestingly, removing methylation from input reference and alternate sequences leads to slightly less agreement between the mean deltas and experimental effect sizes (Spearman ρ = 0.462, signed accuracy = 75.1%), suggesting that ModBERT integrates methylation and genetic variants when capturing individual-specific chromatin accessibility patterns (**Supplementary Figure 4**). In conclusion, these results demonstrate that probabilistic fine-tuning allows ModBERT to generalize to new individuals and new genomic regions, and that its predictions capture the effects of likely causal regulatory variants and methylation on chromatin accessibility.

## DISCUSSION

In this work, we developed ModBERT, a novel DNA LLM that enables incorporation of genetic variants (SNPs) and DNA methylation information in pretraining and in various downstream predictive tasks. ModBERT’s pretrained embeddings already capture genomic elements and exhibit methylation sensitivity, with specific patterns learned in different layers. Using our novel probabilistic fine-tuning framework, ModBERT achieved robust performance gains over ab initio baseline models and its pretrained methylation-ablated version over tasks encompassing transcription factor binding, histone mark peak prediction, and ATAC-seq peak prediction both within and across individuals. ModBERT’s recapitulation of known differential methylation sensitivity across different histone marks and transcription factors demonstrates its understanding of methylation’s role in the regulatory genome. ModBERT’svariant effect prediction performance for caQTLs for both methylated and demethylated sequences highlights its ability to learn associations between methylation levels and genetic variation during fine-tuning.

In terms of limitations, ModBERT was pretrained on only lymphoblastoid cell lines, and given that methylation patterns are cell type-specific^23^, it is unlikely that ModBERT’s learned representations will be applicable to other, more distant cell types. In addition, ONT natively provides phased methylation and genetic variants, which is ignored when providing input sequences into ModBERT during both pretraining and fine-tuning. This prevents the model from learning which variants are in cis with which methylation patterns and obscures allele-specific methylation, genomic imprinting, and the haplotype-dependent effects of variants and methylation on TF binding. Providing phased inputs is a natural extension that we leave to future work. We also note that while robust, the performance gains from incorporating DNA methylation during pretraining were small, suggesting that a model with a larger receptive field may be needed to learn more complex relationships among DNA methylation, genetic variation and gene regulation. Future work could incorporate additional base modifications or epigenomic layers during pretraining to explore this.

## METHODS

### 1KGP-LRSC ONT Data Preprocessing

Haplotagged ONT BAM and VCF files aligned to reference genome GRCh38 were downloaded for 242 lymphoblastoid samples from the 1KGP-LRSC AWS server: https://s3.amazonaws.com/1000g-ont/index.html. These were the total number of samples sequenced with the higher accuracy ONT R10 chemistry available at the time of download. Methylation was quantified at the individual sample level using ONT’s Modkit v0.5.0 software^24^. Specifically, we ran `modkit pileup` on the bam files for just the autosomal chromosomes with the`--cpg, -- ignore h`,and `-- combine-strands` flags to restrict methylation quantification to CpG sites, ignore 5-hydroxymethylcytosine (another base modification present in some of the bam files) by redistributing its probability between canonical C and 5mC, and sum positive and negative strand read counts respectively. Methylation was quantified at the population level by filtered out SNPs with alternate allele frequency less than 2%.

### Input Representation

summing methylation fractions at individual CpG sites across samples and dividing by 242. These average fractions were used to construct probability matrices for pretraining. As for SNP quantification, we first merged all 242 VCF files and subsequently computed allele frequencies using BCFtools^25^. We then We represent a DNA sequence of length L as a 10 x L probabilistic matrix where the last five rows correspond to the bases {A, C, G, T, M} respectively, where “M” represents 5mC. The first five rows correspond to special tokens that are standard for BERT^26^ models: [PAD], [UNK], [CLS], [SEP], [MASK]. For individual donors, single nucleotide variants (SNVs) are one-hot encoded if homozygous or split 50/50 if heterozygous. At CpG sites, the “C” probability is divided into “C” and “M” based on the fraction of the respective modified reads. For population-level data, SNPs and CpG sites are instead encoded using population allele frequencies and donor-averaged methylation frequencies, respectively (see **1KGP-LRSC Data Preprocessing)**. If a CpG site is also a variant, then the entries for “C” and “M” are scaled multiplicatively such that the sum over all five bases remains 1. For all methylation ablation experiments, the probability assigned to “M” is redistributed to the probability assigned to “C”.

### Model Architecture

ModBERT’s architecture is based on MutBERT’s architecture. It takes the probabilistic matrix as input and applies a linear layer to generate learned d-dimensional embeddings, thus each 1-bp locus is represented as a weighted sum of all embeddings. The input embeddings are fed into a 12-layer bidirectional Transformer encoder with a hidden size of 768 and 12 attention heads. To effectively handle variable sequence length during fine-tuning, the model incorporates Rotary Position Embeddings (RoPE)^27^. ModBERT also utilizes Flash Attention, a hardware-aware, memory-efficient implementation of scaled dot-product attention (SDPA)^28^.

### Pretraining

ModBERT was pretrained on 30 million 512-bp length sequences randomly sampled from the human genome overlaid with population-level genetic variability and DNA methylation frequencies derived from Oxford Nanopore technology reads. Sequences were drawn from autosomal chromosomes with unmapped regions omitted. The models were trained utilizing a masked language modeling objective, where 15% of the sequence positions were randomly selected for masking by setting the positions to the [MASK] token. Each selected position is then independently replaced with a one-hot encoded [MASK] vector with probability 0.8. Otherwise it is left unchanged, keeping its original probability distribution. Because the model inputs are probabilistic, the network optimizes a soft DNA cross-entropy loss that computes the negative log-likelihood directly against the soft target distributions rather than standard discrete labels. The weights were optimized using the AdamW^29^ (β1 = 0.9, β2 = 0.999, ε = 10?8) optimizer with a weight decay of 0.01 and a warmup cosine learning rate scheduler with a peak learning rate of 4 × 10^?4^. Gradients were clipped to a global L2 norm of 1.0. Training ran for 120,000 optimizer steps with a batch size of 256 and a gradient accumulation of 4, resulting in ∼4 passes over the 30 million sampled sequences. To assess the impact of 5mC during pretraining, we also pretrained a separate five-letter model with identical architecture (including the “M” row of the probabilistic matrix) and hyperparameters on sequences without 5mC (setting “M” to 0 and distributing its value to “C”). Pretraining was done on a single NVIDIA GH200 Grace Hopper Superchip with fp16 mixed precision and dynamic loss scaling.

### Zero-shot Genomic Element Analysis

Genomic element annotations were derived from the Ensembl^30^ Release 116 GRCh38 GTF file: https://ftp.ensembl.org/pub/release-116/gtf/homo_sapiens/Homo_sapiens.GRCh38.116.gtf.gz. Coding sequence (CDS), 5’ untranslated region (5’ UTR), 3’ untranslated region (3’ UTR) were directly pulled from the annotation file. Introns were defined as gene regions minus exons. Intergenic regions were obtained from subtracting gene regions from the genome. A strict disjoint hierarchy (CDS > 5’UTR > 3’UTR > Intron > Intergenic) was then applied by sequentially subtracting higher priority elements from lower priority ones to eliminate overlapping elements. We then sampled 5,000 510-bp sequences (in probability matrix form) from each genomic element on autosomal chromosomes, ensuring that each sequence lay entirely within an element. SNP and methylation frequencies were derived from the 1KGP-LRSC cohort. These matrices were subsequently fed into the pretrained ModBERT model to obtain 768-dimensional sequence embeddings from the output of each of the 12 layers via mean pooling of the given layer’s hidden state token embeddings. Lastly, UMAP^31^ was applied to each layer’s embeddings using cosine distance as a metric, 15 neighbors, and a minimum distance = 0.1.

### Zero-shot Simulated Methylation Gradient

We sampled 1,000 510-bp length sequences from autosomal chromosomes, each with exactly 30 CpG sites. All CpG sites were uniformly tuned to 0%, 20%, 40%, 60%, 80%, and 100% within each sequence. Each of these perturbed sequences was fed into the pretrained ModBERT model to obtain layer-specific sequence embeddings in the same manner as the genomic element analysis. A layer-specific longitudinal analysis for each sequence was then performed by computing the cosine similarity of each non-zero methylated sequence’s embedding with respect to its unmethylated sequence embedding.

### GM12878 ONT Data Preprocessing

ONT GRCh38-aligned bam files were downloaded for two flowcells (PAW79146 and PAW81754) from the Genome in a Bottle January 2025 release^32^: s3://ont-open-data/giab_2025.01/basecalling/sup/HG001. These data were also generated using R10 technology as with the 1KGP-LRSC cohort. We ran modkit on each flowcell’s bam file separately using the same flags as the 1KGP-LRSC cohort. We then computed the Pearson correlation between CpG site methylation percentages of the two flowcells per-chromosome. Correlation values ranged from 0.96 to 0.98, indicating minimal batch effect and justifying the merging of the bam files. We subsequently merged the bam files using SAMtools 1.23.1 and called methylation percentages on the merged file, which were then used for all downstream fine-tuning. VCF files were downloaded for the same flow cells (s3://ont-open-data/giab_2025.01/analysis/wf-human-variation/sup/HG001/${FLOWCELL}/output/SAMPLE.wf_snp.vcf.gz) and merged using BCFtools.

### Fine-tuning

ModBERT was finetuned on several sequence classification tasks. For all fine-tuning tasks, each sample-specific input sequence was represented as a probabilistic matrix as in the pretraining with all model weights updated during training (full-parameter fine-tuning). A classification head consisting of Each task is a binary prediction to determine whether a fixed-length sequence overlaps with a “peak” or not. Positive sequences were constructed by extracting fixed-length windows from the centers of peaks. GC-content matched negative sequences were extracted by random sampling of the reference genome without overlapping any peaks at a 1:1 ratio with positive instances, ensuring balanced training, validation, and test sets.

For all tasks, when training and testing were done on GM12878, chromosomes 5, 10, 14, 18, 20, 22 were left out for testing with validation performed on chromosomes 6 and 21. The same chromosomes were used for validation and testing for the unseen peaks and unseen samples and peaks tasks. To assess the impact of pretraining, we also trained the following baseline models ab initio: Long Short Term Memory (LSTM), Recurrent Neural Network (RNN), Gated Recurrent Unit (GRU), and Convolutional Neural Network (CNN). The LSTM, GRU, and RNN models are all bi-directional and composed of two layers with hidden dimension = 128. The per-position representations are collapsed into a 256-dimensional vector via max pooling that is then fed into a two-layer classification head with dimensions (256, 64, 2). The CNN model consists of two convolutional layers (128, 256), each with kernel size 7, stride 1, and zero-padding of 3, which preserves sequence length. As with the recurrent models, the output of these layers was collapsed via max pooling and fed into a two-layer classifier with dimensions (256, 64, 2). All baseline and ModBERT models were trained with a learning rate of 3e-5 and batch size 32 for 10 epochs with early stopping if accuracy did not improve after 3 epochs on the validation set. Fine-tuning was done on NVIDIA A40 GPUs.

### Transcription Factor Methylation Sensitivity Scores

TF methylation sensitivity scores were derived from the methylation-aware position weight matrix models of Grau et al. (2023), which were obtained from https://doi.org/10.5281/zenodo.3723984. In these models the DNA alphabet is extended with two letters: M, for a methylated cytosine in a CpG context, and H, for the guanine opposite a methylated cytosine. The models were trained on 1000-bp, methylation-aware sequences (in which methylation calls from whole-genome bisulfite sequencing were discretised) centered on GM12878 ENCODE ChIP-seq peaks. Each TF has ten models, each one belonging to a different cross-validation fold. We utilized the same GM12878 ENCODE ChIP-seq data as Grau et al. for binding site prediction. For each of the cross-validation folds, the top-ranked motif was used to predict one binding site per peak (the best scoring window on either strand), and per-position methylation sensitivity profiles were computed as described by Grau et al. Specifically, at each motif position *i*, the sensitivity is *s*_*i*_ = *f*_*i*_ · *δ*_*i*_, where *f*_*i*_ is the fraction of predicted sites with a CpG at positions *i, i*+1, and *δ*_*i*_ is the change in model score when that CpG is converted to MpH. Sites were predicted from all peaks rather than from each fold’s training partition (which contains 90% of the peaks) because the training splits were not available to us. Following Grau et al., each fold’s profile was summarized by its minimum and maximum. For each TF, we took the medians of the per-fold minima and maxima separately and used whichever had the larger absolute value as the TF’s methylation sensitivity score. Negative scores indicate that the model disfavors methylated binding sites. TFs were compared to ModBERT’s predicted changes in binding probability only if their motif contained a prominent CpG, defined as a position at which more than 25% of predicted sites contained a CpG (median across folds), leaving the 44 TFs in **Figure 4**.

### Transcription Factor Motif Attention Analysis

To determine whether the fine-tuned models preferentially attend to TF binding motifs, we first annotated motif occurrences in the held-out test set of each TF ChIP-seq dataset. To obtain a nucleotide sequence for motif scanning, the 5mC probability was added to cytosine, and each position was assigned the base with the highest resulting probability. Motif occurrences were identified with FIMO using position weight matrices from the HOCOMOCO v14 core collection. FIMO was run with default parameters, reporting matches on both strands with a p-value threshold of 10?4. For each dataset, we loaded the fine-tuned ModBERT model. Only positive (ChIP-seq peak) test sequences containing at least one FIMO motif occurrence were analysed. The full probabilistic matrix, including methylation and SNV information, was passed through the model to obtain attention matrices. For each sequence, attention matrices were averaged across all layers and all attention heads to produce a single L × L matrix (L = 101), where rows correspond to query positions and columns to key positions. A per-position attention importance score was then computed as the mean attention each position receives, averaged over all query positions. Because each row of the attention matrix sums to one, the expected score under uniform attention is 1/L ≈ 0.0099. A binary mask was constructed from the FIMO start and stop coordinates, marking the positions spanned by the motif occurrence. For each sequence, we computed the mean importance score over motif positions and over all remaining (non-motif) positions. These values were then averaged across all sequences within a dataset, producing one mean motif attention score and one mean non-motif attention score per TF.

### ATAC-seq Data Preprocessing

Kumasaka et al. (2019) CRAM files for 27 1000 Genomes samples that were present in the 1KGP-LRSC cohort were downloaded from the European Nucleotide Archive^33^ under accession ERP110508 (**Supplementary Table 3**). Because these files were originally aligned to the GRCh37 reference genome, we first converted them to FASTQ files using samtools fastq. Alignment to reference genome GRCh38 and peak calling were then performed using our in-house ATAC-seq processing pipeline: https://github.com/ay-lab/ATACProc.

### In Silico Mutagenesis of caQTLs

We used caQTLs defined in Kumasaka et al. (2019). Each caQTL-peak pair is assigned an effect size β and a probability of the caQTL being causal for the peak (Causal Prob). We lifted their population-level ATAC-seq peaks (the caQTL targets) and caQTLs from GRCh37 to GRCh38 using CrossMap^34^ and intersected peaks with each sample’s 201-bp positive sequence. We kept lead SNPs falling inside a positive sequence from unseen samples and unseen chromosomes. In each sequence we then set the caQTL to either the reference or the alternative allele, leaving the rest of the sample’s sequence and methylation unchanged and computed a Delta = P(accessible | ALT) ™ P(accessible | REF) using the fine-tuned unseen samples and peaks by the ModBERT model. Delta was averaged across all samples that contained a positive sequence with the caQTL to give one score per caQTL–peak pair as shown in **Figure 5e**. We then evaluated this mean delta using a signed accuracy, i.e., the fraction of caQTL-peak pairs for which sign(Mean Delta) = sign(β). Because a lead variant may only tag the causal variant through linkage disequilibrium, we also stratified pairs by Causal Prob where we defined high as > 0.999 and low as < 0.01. We also repeated the ISM by setting methylation to zero for all input sequences.

## Acknowledgements

Research supported by the NVIDIA Academic Grant Program using 8 A100 GPUs and by the R35-GM128938 grant from National Institutes of Health (NIH) to F.A.

## Code Availability

https://github.com/ay-lab/ModBERT

## Notes

### Competing Interest Statement

The authors have declared no competing interest.

