## Supplementary Figure 1b for "ModBERT: a DNA modification and genetic variant-aware DNA large language model"

| Histone Mark | File Accession |
|---|---|
| H2AFZ | ENCFF377OJG |
| H3K27ac | ENCFF023LTU |
| H3K27me3 | ENCFF695ETB |
| H3K36me3 | ENCFF475QVQ |
| H3K4me1 | ENCFF321BVG |
| H3K4me2 | ENCFF283LNH |
| H3K4me3 | ENCFF998CEU |
| H3K79me2 | ENCFF131ZGZ |
| H3K9ac | ENCFF981JOU |
| H3K9me3 | ENCFF725UFY |
| H4K20me1 | ENCFF469QYW |

**Table 1**: Histone mark ChIP-seq ENCODE bed file accessions

| TF | File Accession |
|---|---|
| ARNT | ENCFF596RIT |
| ATF2 | ENCFF213LSA |
| ATF3 | ENCFF882AEU |

| | |
|---|---|
| ATF7 | ENCFF277FJJ |
| BACH1 | ENCFF631RZH |
| BATF | ENCFF728KFD |
| CEBPB | ENCFF864EON |
| CREB1 | ENCFF053UZX |
| CREM | ENCFF294IGP |
| CTCF | ENCFF796WRU |
| CUX1 | ENCFF451AII |
| E2F4 | ENCFF744QAC |
| E2F8 | ENCFF961ZFW |
| E4F1 | ENCFF855PXT |
| EGR1 | ENCFF612EIU |
| ELF1 | ENCFF479HJF |
| ELK1 | ENCFF164MPE |
| ETS1 | ENCFF568AZT |
| ETV6 | ENCFF151UJT |
| FOS | ENCFF571DGT |
| FOXK2 | ENCFF639XYN |

| | |
|---|---|
| FOXM1 | ENCFF549GKZ |
| GABPA | ENCFF093KLR |
| HSF1 | ENCFF663ODL |
| IRF3 | ENCFF887WLX |
| IRF4 | ENCFF113VGD |
| IRF5 | ENCFF478SRO |
| JUNB | ENCFF912OPT |
| JUND | ENCFF134BQO |
| KLF5 | ENCFF992IBT |
| MAFK | ENCFF436SJS |
| MAX | ENCFF361EVH |
| MAZ | ENCFF250FJC |
| MEF2A | ENCFF826GQU |
| MEF2B | ENCFF884QQW |
| MEF2C | ENCFF238UKB |
| MXI1 | ENCFF376AEL |
| MYB | ENCFF705CGM |
| MYC | ENCFF214XPD |

| | |
|---|---|
| NFIC | ENCFF980SDE |
| NFYA | ENCFF937QTP |
| NFYB | ENCFF156MUM |
| NR2C1 | ENCFF626EEU |
| NR2C2 | ENCFF782KRV |
| NRF1 | ENCFF864EBU |
| PAX5 | ENCFF192WNJ |
| PAX8 | ENCFF934NVD |
| PBX3 | ENCFF402DQD |
| RELB | ENCFF968MBN |
| REST | ENCFF262MRD |
| RFX5 | ENCFF601WHE |
| RUNX3 | ENCFF346JDW |
| RXRA | ENCFF063KLF |
| SMAD5 | ENCFF485GWN |
| SP1 | ENCFF038AVV |
| SPI1 | ENCFF492ZRZ |
| SRF | ENCFF432PLY |

| | |
|---|---|
| STAT1 | ENCFF973ACG |
| STAT3 | ENCFF029PAA |
| TAF1 | ENCFF502IPK |
| TBP | ENCFF099PYR |
| TBX21 | ENCFF170IZO |
| TCF7 | ENCFF900GTF |
| USF1 | ENCFF879TPT |
| USF2 | ENCFF196JQW |
| ZBED1 | ENCFF753UAX |
| ZBTB40 | ENCFF171FMR |
| ZEB1 | ENCFF703XCL |
| ZNF24 | ENCFF664TYB |

**Table 2**: TF ChIP-seq ENCODE bed file accessions

| 1000 Genomes Sample | CRAM File ENA Directory |
|---|---|
| HG00097 | https://ftp.sra.ebi.ac.uk/vol1/analysis/ERZ683/ERZ683886 |
| HG00102 | https://ftp.sra.ebi.ac.uk/vol1/analysis/ERZ683/ERZ683869 |
| HG00109 | https://ftp.sra.ebi.ac.uk/vol1/analysis/ERZ683/ERZ683927 |

| | |
|---|---|
| HG00112 | https://ftp.sra.ebi.ac.uk/vol1/analysis/ERZ683/ERZ683840 |
| HG00113 | https://ftp.sra.ebi.ac.uk/vol1/analysis/ERZ683/ERZ683847 |
| HG00115 | https://ftp.sra.ebi.ac.uk/vol1/analysis/ERZ683/ERZ683858 |
| HG00116 | https://ftp.sra.ebi.ac.uk/vol1/analysis/ERZ683/ERZ683935 |
| HG00119 | https://ftp.sra.ebi.ac.uk/vol1/analysis/ERZ683/ERZ683846 |
| HG00142 | https://ftp.sra.ebi.ac.uk/vol1/analysis/ERZ683/ERZ683882 |
| HG00143 | https://ftp.sra.ebi.ac.uk/vol1/analysis/ERZ683/ERZ683884 |
| HG00145 | https://ftp.sra.ebi.ac.uk/vol1/analysis/ERZ683/ERZ683903 |
| HG00154 | https://ftp.sra.ebi.ac.uk/vol1/analysis/ERZ683/ERZ683885 |
| HG00155 | https://ftp.sra.ebi.ac.uk/vol1/analysis/ERZ683/ERZ683895 |
| HG00160 | https://ftp.sra.ebi.ac.uk/vol1/analysis/ERZ683/ERZ683877 |
| HG00231 | https://ftp.sra.ebi.ac.uk/vol1/analysis/ERZ683/ERZ683851 |
| HG00234 | https://ftp.sra.ebi.ac.uk/vol1/analysis/ERZ683/ERZ683902 |
| HG00236 | https://ftp.sra.ebi.ac.uk/vol1/analysis/ERZ683/ERZ683892 |
| HG00237 | https://ftp.sra.ebi.ac.uk/vol1/analysis/ERZ683/ERZ683844 |
| HG00246 | https://ftp.sra.ebi.ac.uk/vol1/analysis/ERZ683/ERZ683928 |
| HG00250 | https://ftp.sra.ebi.ac.uk/vol1/analysis/ERZ683/ERZ683934 |
| HG00253 | https://ftp.sra.ebi.ac.uk/vol1/analysis/ERZ683/ERZ683888 |

| | |
|---|---|
| HG00256 | https://ftp.sra.ebi.ac.uk/vol1/analysis/ERZ683/ERZ683878 |
| HG00259 | https://ftp.sra.ebi.ac.uk/vol1/analysis/ERZ683/ERZ683937 |
| HG00262 | https://ftp.sra.ebi.ac.uk/vol1/analysis/ERZ683/ERZ683849 |
| HG00263 | https://ftp.sra.ebi.ac.uk/vol1/analysis/ERZ683/ERZ683889 |
| HG00264 | https://ftp.sra.ebi.ac.uk/vol1/analysis/ERZ683/ERZ683863 |
| HG01334 | https://ftp.sra.ebi.ac.uk/vol1/analysis/ERZ683/ERZ683931 |

**Table 3**: ATAC-seq samples from Kumasaka et al., 2019. European Nucleotide Archive (ENA) source directories provided for the respective CRAM files.

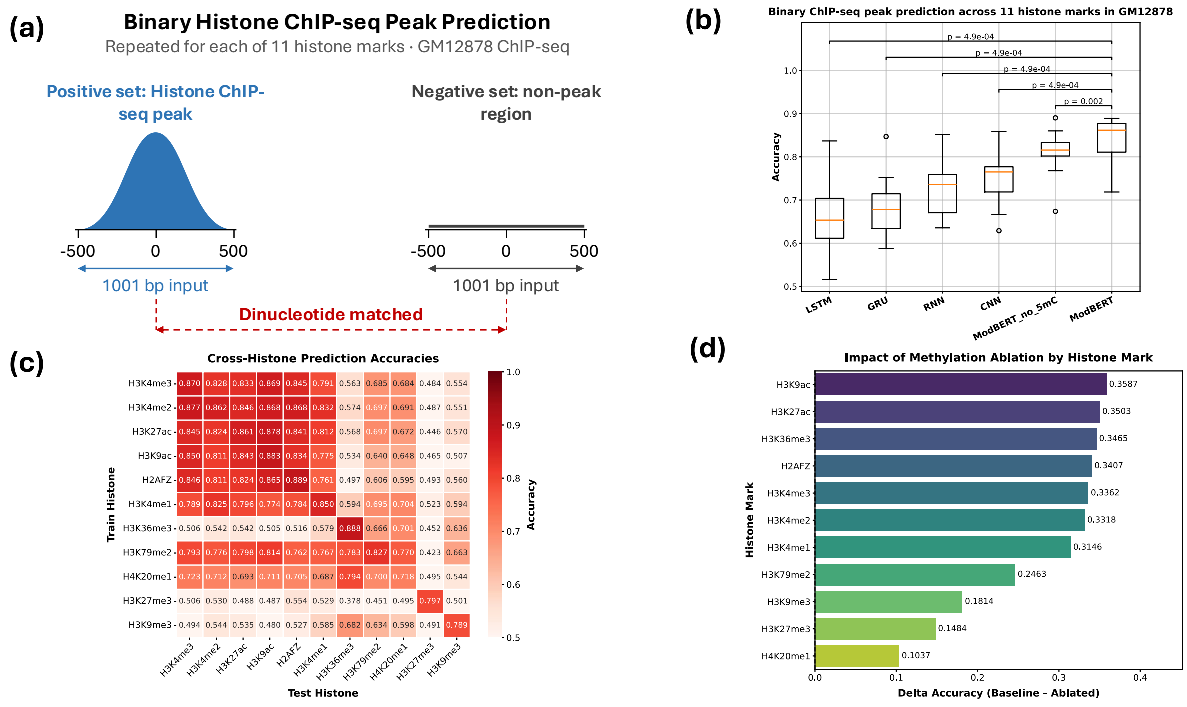


**Supplementary Figure 1**: **Binary histone ChIP-seq peak prediction across 11 histone marks in GM12878.** Same analysis as Main Figure 3, but with dinucleotide-matched negatives in lieu of GC content matched negatives.

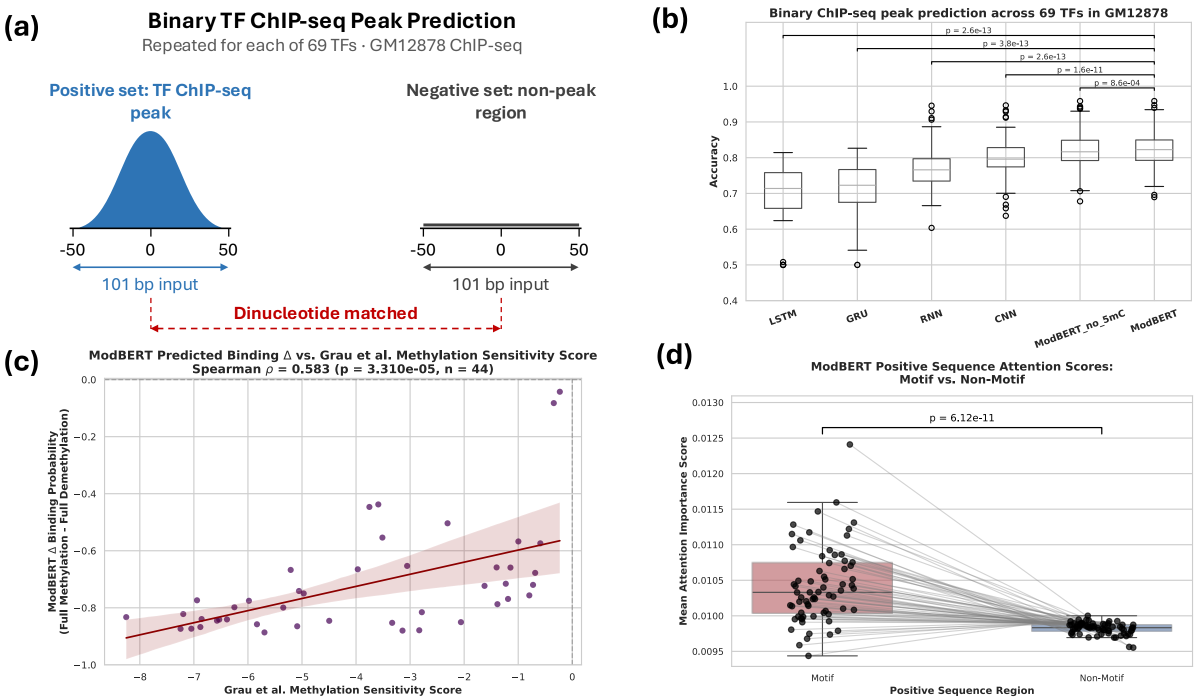


**Supplementary Figure 2**: **Binary TF ChIP-seq peak prediction across 69 TFs.** Same analysis as Main Figure 4, but with dinucleotide-matched negatives in lieu of GC content matched negatives.

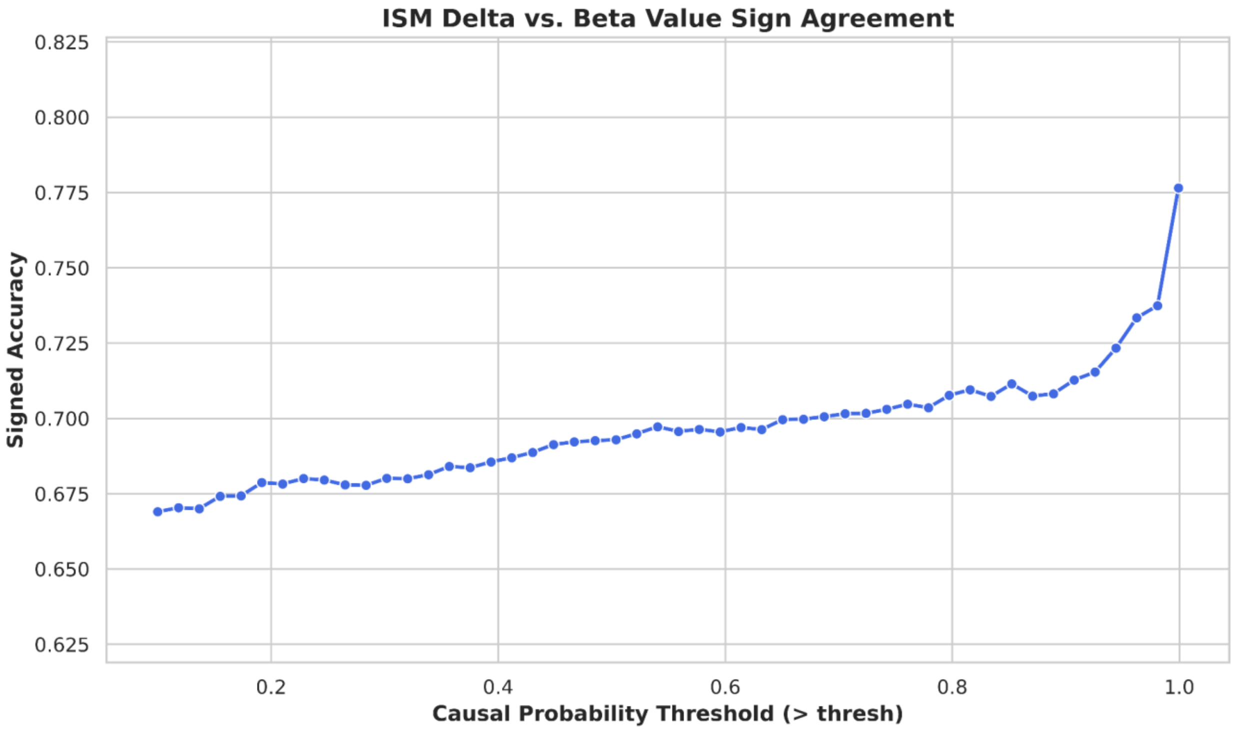

**Supplementary Figure 3**: ModBERT signed accuracy from in silico mutagenesis of caQTLs as a function of the causal probability threshold for filtering them. The thresholds are 50 evenly spaced points between 0.1 and 0.999.

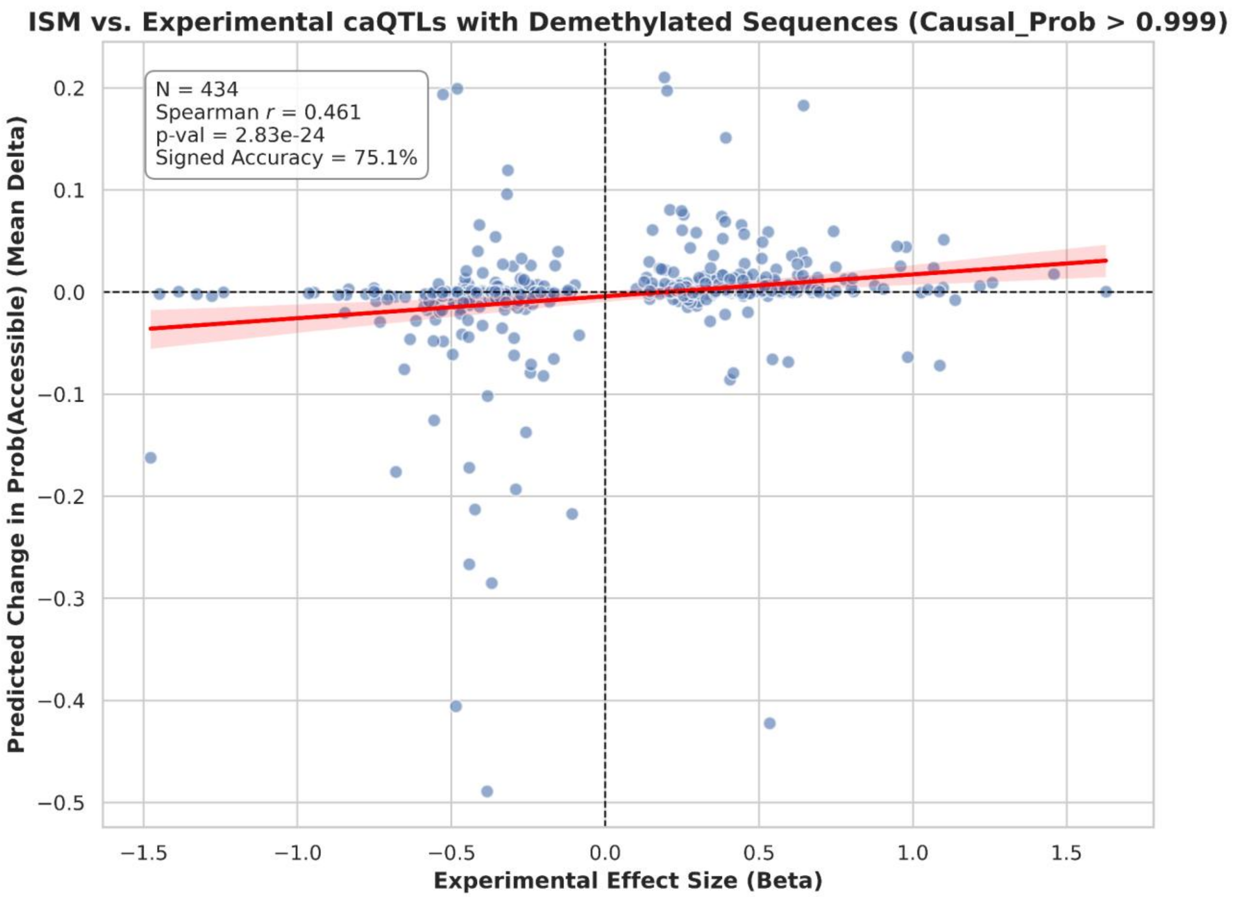


**Supplementary Figure 4**: In silico mutagenesis of the same set of caQTLs as **Figure 5**, but with methylation in all CpG sites set to 0.
